# High-dose pyruvate treatment alters skeletal muscle differentiation and expression of inflammation-related genes

**DOI:** 10.1101/2020.09.21.305847

**Authors:** Kazuya Hasegawa, Yuya Yamaguchi, Yutthana Pengjam

**Author notes:** Address for correspondence: Kazuya Hasegawa, M.D., Ph.D., Faculty of Nutritional Sciences, Morioka University, 808 Sunakomi, Takizawa city, Iwate 020-0694, Japan.

## Abstract

Pyruvic acid therapy is used for various diseases, but the therapeutic effect decreases at high doses. The molecular mechanism of high-dose pyruvate is not well understood. The purpose of this study was to identify the effects of high dose pyruvate addition on skeletal muscle using C2C12. The gene expression profile for the GSE5497 dataset was taken from the Gene Expression Omnibus database. GEO2R was used to identify specifically expressed genes (DEGs). Functional analysis and pathway enrichment analysis of DEG were performed using the DAVID database. The protein-protein interaction (PPI) network was built in the STRING database and visualized using Cytoscape. GO analysis showed that up-regulated DEG was primarily involved in angiogenesis, cell adhesion, and inflammatory response. We also showed that down-regulated DEG is involved in the regulation of muscle contraction, skeletal muscle fiber development. In addition, the upregulated KEGG pathway of DEG included Rheumatoid arthritis, Chemokine signaling pathway, and Cytokine-cytokine receptor interaction. Downregulated DEG included Calcium signaling pathway, hypertrophic cardiomyopathy (HCM), Dilated cardiomyopathy, Neuroactive ligand-receptor interaction, and Cardiac muscle contraction. Further, analysis of two modules selected from the PPI network showed that high-dose pyruvate exposure to C2C12 was primarily associated with muscle contraction, muscle organ morphogenesis, leukocyte chemotaxis, and chemokine activity. In conclusion, High-dose pyruvate treatment of C2C12 was found to be associated with an increased inflammatory response and decreased skeletal muscle formation. However, further studies are still needed to verify the function of these molecules at high doses of pyruvate.

## INTRODUCTION

Pyruvic acid is an important molecule in many aspects of eukaryotic and human metabolism. Pyruvic acid is the final product of glycolysis and is ultimately destined for transport to mitochondria as a master fuel to support the carbon flux of the citric acid cycle (1,2). In mitochondria, pyruvate promotes ATP production through multiple biosynthetic pathways that intersect the oxidative phosphorylation and citric acid cycle (1,2). Administration of pyruvic acid suppresses inflammation and oxidative stress (3-6). Currently, pyruvate therapy is used for a variety of diseases (7-10). However, high doses of pyruvate diminish the therapeutic effect (9). In contrast to low doses, the molecular mechanism of high doses of pyruvate has not been clarified.

In this study, we used the gene expression profile of high-dose pyruvate-treated C2C12 skeletal myocytes from the GEO database to identify differentially expressed genes (DEGs). Next, Gene Ontology (GO) enhancement analysis and Kyoto Encyclopedia of Genes and Genomes (KEGG) pathway enhancement analysis were further performed to analyze the major biological functions of co-modulated DEG. Next, we constructed a protein-protein interaction (PPI) network and identified hub genes and modules associated with GC by STRING and Cytoscape. The results of current bioinformatics analysis may help elucidate the molecular mechanism of high-dose pyruvate by identifying the major genes and pathways that contribute to the phenotype.

## MATERIALS AND METHODS

### Microarray data

The GEO database (https://www.ncbi.nlm.nih.gov/geo/) stores the original records and curated datasets provided by the submitter. The gene expression dataset GSE5497 was obtained from the GEO database. The dataset GSE5497 contained three C2C12 samples that received 50 mM pyruvate and three C2C12 (controls) samples that did not receive pyruvate. This dataset is a platform Generated by Wilson L et al. Using the GPL1261 [Mouse430_2] Affymetrix Mouse Genome 430 2.0 Array (11). This dataset contains a total of 6 samples and consists of a control (3 samples) and a high dose of pyruvate (3 samples) of C2C12. Baseline characteristic data for the sample was previously published by Wilson L et al (11). Therefore, a total of 6 chips could be used for subsequent analysis.

### GO and KEGG pathway enrichment analysis

A database for annotation, visualization and integrated discovery (DAVID 6.8, http://david.ncifcrf.gov/summary.jsp) is used to systematically extract biological implications from a large list of genes and proteins. GO enrichment and KEGG pathway analysis were performed using the DAVID online tool to analyze the screened DEG at the functional level. P <0.05 was set as the cutoff criterion.

### PPI network analysis

The PPI network was used to further evaluate the functional interactions between the DEGs. DEGs are mapped using a search tool (STRING; version 11.0; http://string-db.org/cgi/input.pl) for searching for interacting genes and have a PPI binding score of 0.4 or higher. Only action pairs were selected as important. The PPI network was then built using Cytoscape software (https://cytoscape.org/; version 3.6.1). The top 10 required nodes ranked in each DEG network were calculated using the Cytoscape plugin CytoHubba.

### Module analysis

We used MCODE to identify tightly connected areas of Cytoscape’s PPI network. I applied it to the screen module of the PPI network with the following parameters: degree cutoff = 10, node score cutoff = 0.2, k-core = 2, max depth = 100. In addition, the function and pathway of DEG for each module was evaluated using ClueGo and P <0.05 was considered significant.

## RESULTS

### Mutual comparative evaluation of microarray data

Expression values were investigated via box plot analysis for logical comparison of defined groups. Box plot analysis shows that the pyruvate-treated sample and the control gene expression profile are consistent (Fig. 1). Therefore, the samples are statistically comparable.

**Figure 1.**
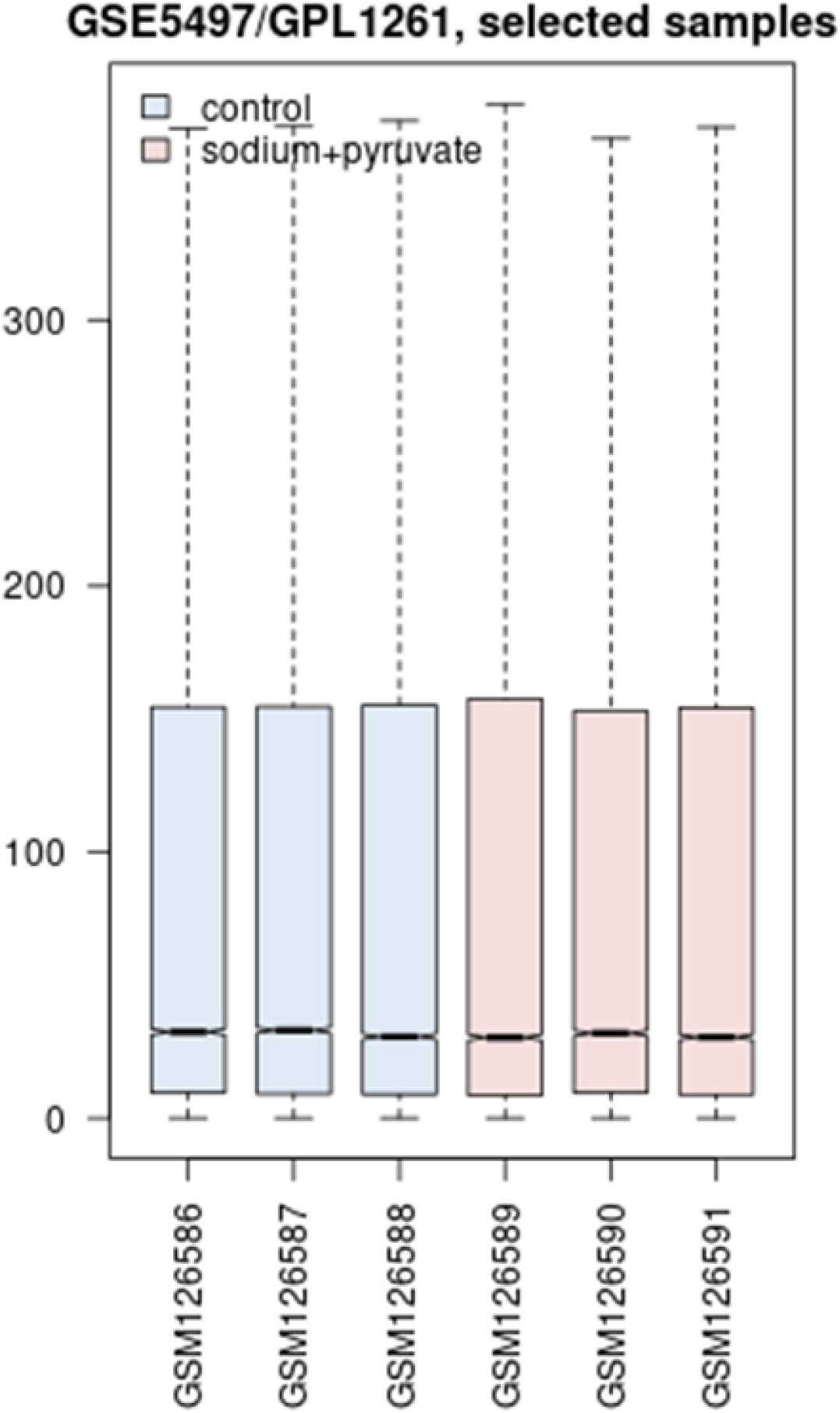
Boxplot indication of median-centered sample of control and pyruvate (50 mM).

### DEG screening

The gene expression dataset GSE5497 was downloaded from the GEO database. The DEG between pyruvate administration and control was determined using the GEO2R tool. Total of 377 DEGs were selected using the cutoff point of P < 0.05 and |log2FC| >1, which included 178 upregulated and 199 downregulated DEGs. Table 1 shows the top 10 upregulated and downregulated genes of pyruvate and controls.

**Table 1.**
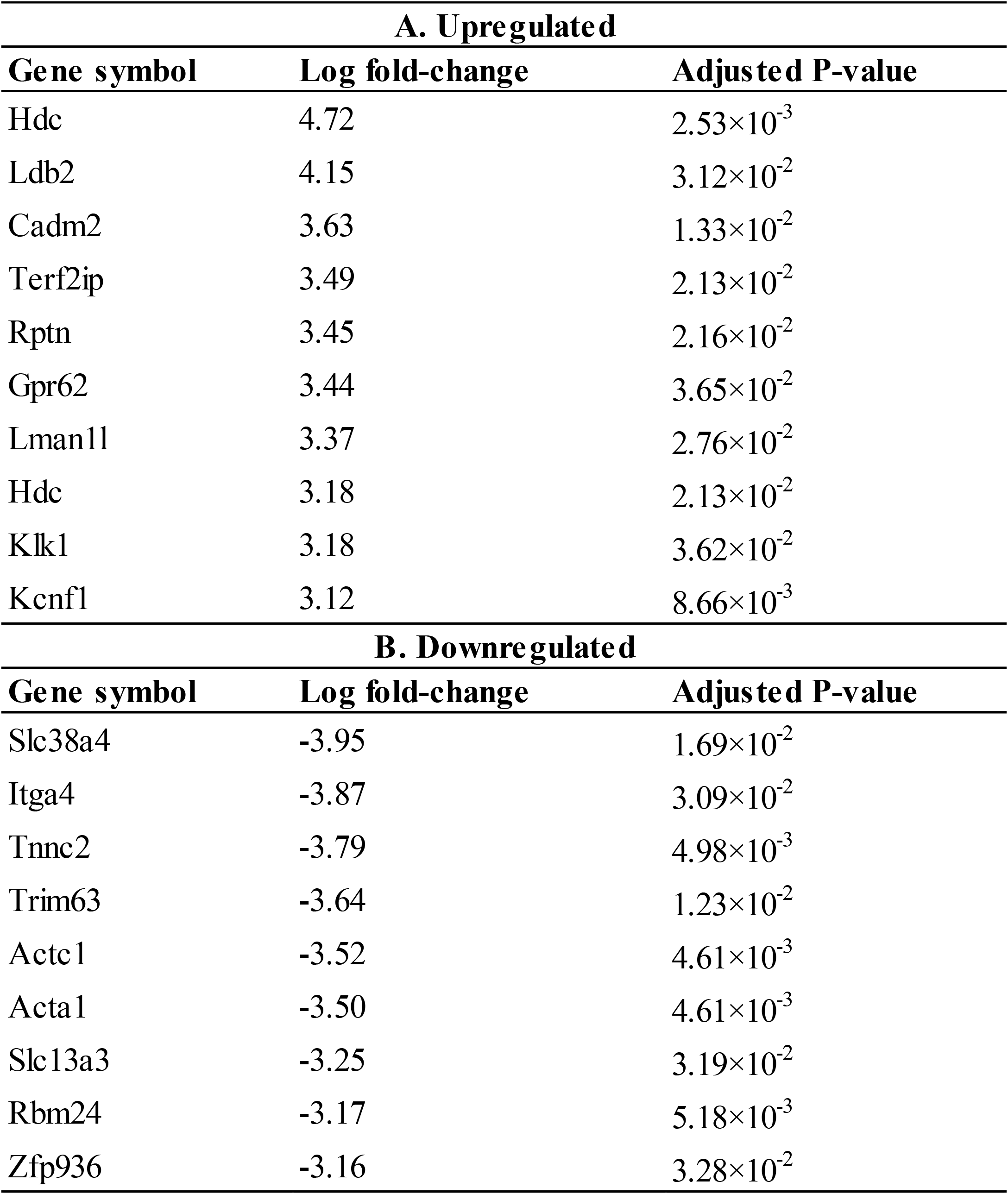
Top 10 upregulated and downregulated genes.

### GO and KEGG pathway enrichment analyses

Functional annotations and pathway analyzes, including GO and KEGG analyzes, were performed using DAVID. Table 2 shows the results of GO analytical biological processes (BP) and KEGG. The genes for which DEG BP was upregulated were mainly rich in angiogenesis, cell adhesion, and inflammatory response. The down-regulated genes were predominantly rich in regulation of muscle contraction and skeletal muscle fiber development.

**Table 2.** Top 5 GO and KEGG pathway of Differentially Expressed Genes after pyruvic acid administration.

| A. Upregulated |  |  |  |  |
| --- | --- | --- | --- | --- |
| Category | Term | Function | Count | P-value |
| Biological process | GO:0001525 | angiogenesis | 12 | $1.81 \times 10^{-6}$ |
| | GO:0007155 | cell adhesion | 16 | $3.47 \times 10^{-6}$ |
| | GO:0006954 | inflammatory response | 11 | $2.62 \times 10^{-4}$ |
| | GO:0070374 | positive regulation of ERK1 and ERK2 cascade | 8 | $5.31 \times 10^{-4}$ |
| | GO:0007157 | heterophilic cell-cell adhesion via plasma membrane cell adhesion molecules | 5 | $6.09 \times 10^{-4}$ |
| KGEE pathway | mmu05323 | Rheumatoid arthritis | 5 | $4.96 \times 10^{-3}$ |
| | mmu04062 | Chemokine signaling pathway | 7 | $6.01 \times 10^{-3}$ |
| | mmu04977 | Vitamin digestion and absorption | 3 | $1.46 \times 10^{-2}$ |
| | mmu04060 | Cytokine-cytokine receptor interaction | 7 | $1.60 \times 10^{-2}$ |
| | mmu04015 | Rap1 signaling pathway | 6 | $3.41 \times 10^{-2}$ |
| B. Downregulated |  |  |  |  |
| Category | Term | Function | Count | P-value |
| Biological process | GO:0006937 | regulation of muscle contraction | 10 | $8.44 \times 10^{-14}$ |
| | GO:0003009 | skeletal muscle contraction | 10 | $3.16 \times 10^{-13}$ |
| | GO:0060048 | cardiac muscle contraction | 10 | $1.46 \times 10^{-10}$ |
| | GO:0006936 | muscle contraction | 7 | $2.96 \times 10^{-6}$ |
| | GO:0048741 | skeletal muscle fiber development | 5 | $6.20 \times 10^{-5}$ |
| KGEE pathway | mmu04020 | Calcium signaling pathway | 7 | $2.04 \times 10^{-3}$ |
| | mmu05410 | Hypertrophic cardiomyopathy (HCM) | 5 | $2.70 \times 10^{-3}$ |
| | mmu05414 | Dilated cardiomyopathy | 5 | $3.23 \times 10^{-3}$ |
| | mmu04080 | Neuroactive ligand-receptor interaction | 7 | $1.83 \times 10^{-2}$ |
| | mmu04260 | Cardiac muscle contraction | 4 | $1.92 \times 10^{-2}$ |

For enrichment analysis of the KEGG pathway, upregulated genes were primarily involved in Rheumatoid arthritis, Chemokine signaling pathway, Vitamin digestion and absorption, Cytokine-cytokine receptor interaction, and Rap1 signaling pathway. The down-regulated genes were mainly involved in the Calcium signaling pathway, hypertrophic cardiomyopathy (HCM), Dilated cardiomyopathy, Neuroactive ligand-receptor interaction, and Cardiac muscle contraction.

### PPI network construction, module and hub gene identification

The PPI network was built by STRING (a database of known and predicted protein interactions), a search tool for searching for interacting genes. The nodes of the PPI network represent genes, and the edges between the nodes represent the interactions between the genes. The network consisted of 298 nodes and 707 edges (Fig. 2). The PPI network then used the Cytoscape plugin CytoHubba to assess the connectivity of the PPI network and identify the top 10 genes (Fig. 3). Titin (Ttn), Myosin light chain 1/3, skeletal muscle isoform (Myl1), Actin, alpha skeletal muscle (Acta1), Troponin T fast skeletal muscle (Tnnt3), Troponin C skeletal muscle (Tnnc2), Troponin T cardiac muscle (Tnnt2), Myosin light chain phosphorylatable fast skeletal muscle (Mylpf), Troponin I fast skeletal muscle (Tnni2), Actin cytoplasmic 1 (Actb), Myogenin (Myog).

**Figure 2.**
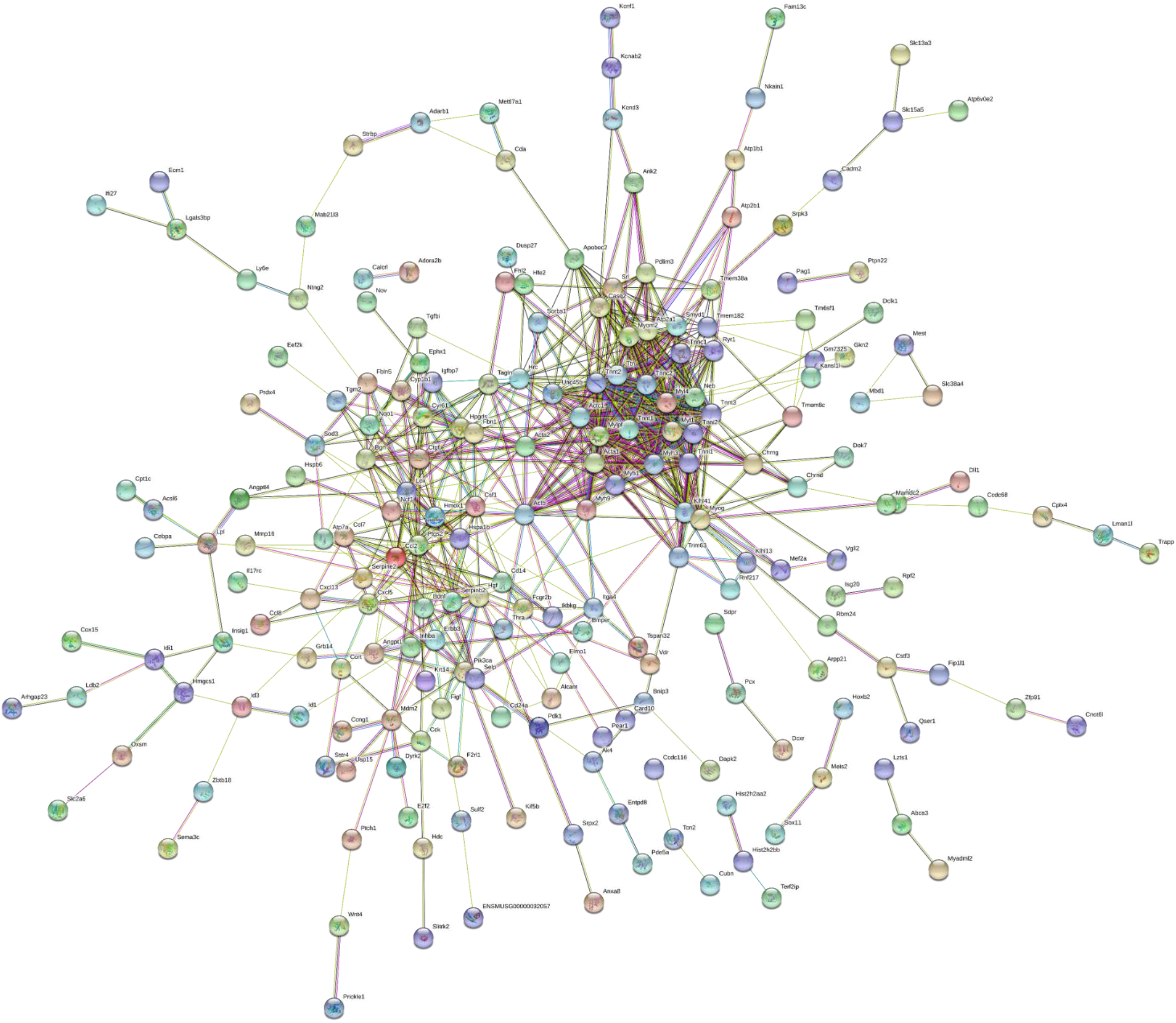
PPI (protein–protein interaction) network of differentially expressed genes. Line thickness indicates the strength of data support.

**Figure 3.**
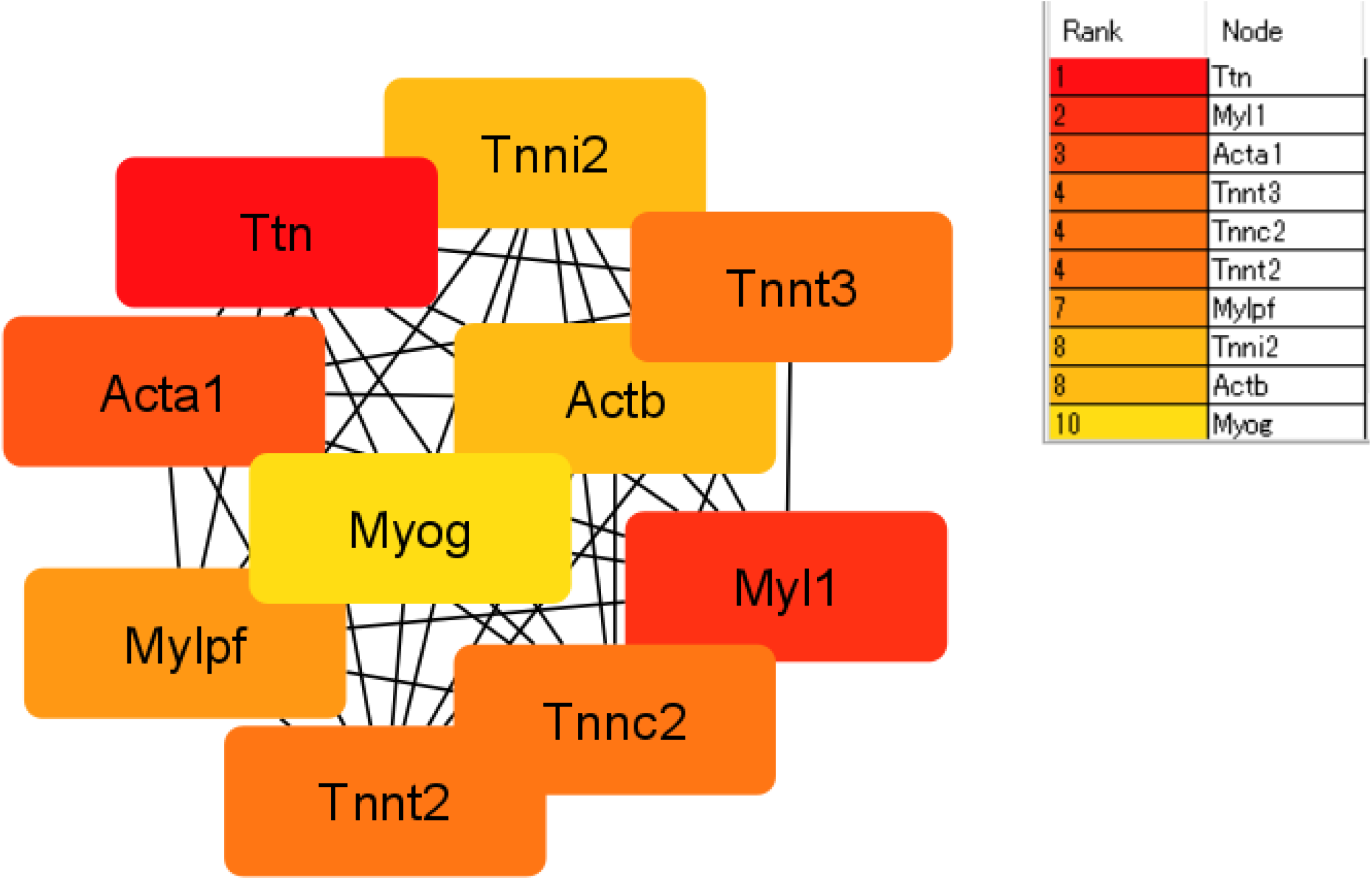
The top 10 genes the highest degree of interaction in the PPI network.

An important module was detected in the PPI network using the MCODE plugin. Ten modules were identified and the top two important modules were selected for subsequent analysis. Module 1 was composed of 20 genes and Module 2 was composed of 12 genes. GO analysis showed that module 1 was primarily associated with muscle contraction and muscle organ morphogenesis, while module 2 was primarily associated with leukocyte chemotaxis, chemotaxis activity(Fig. 4,5).

**Figure 4.**
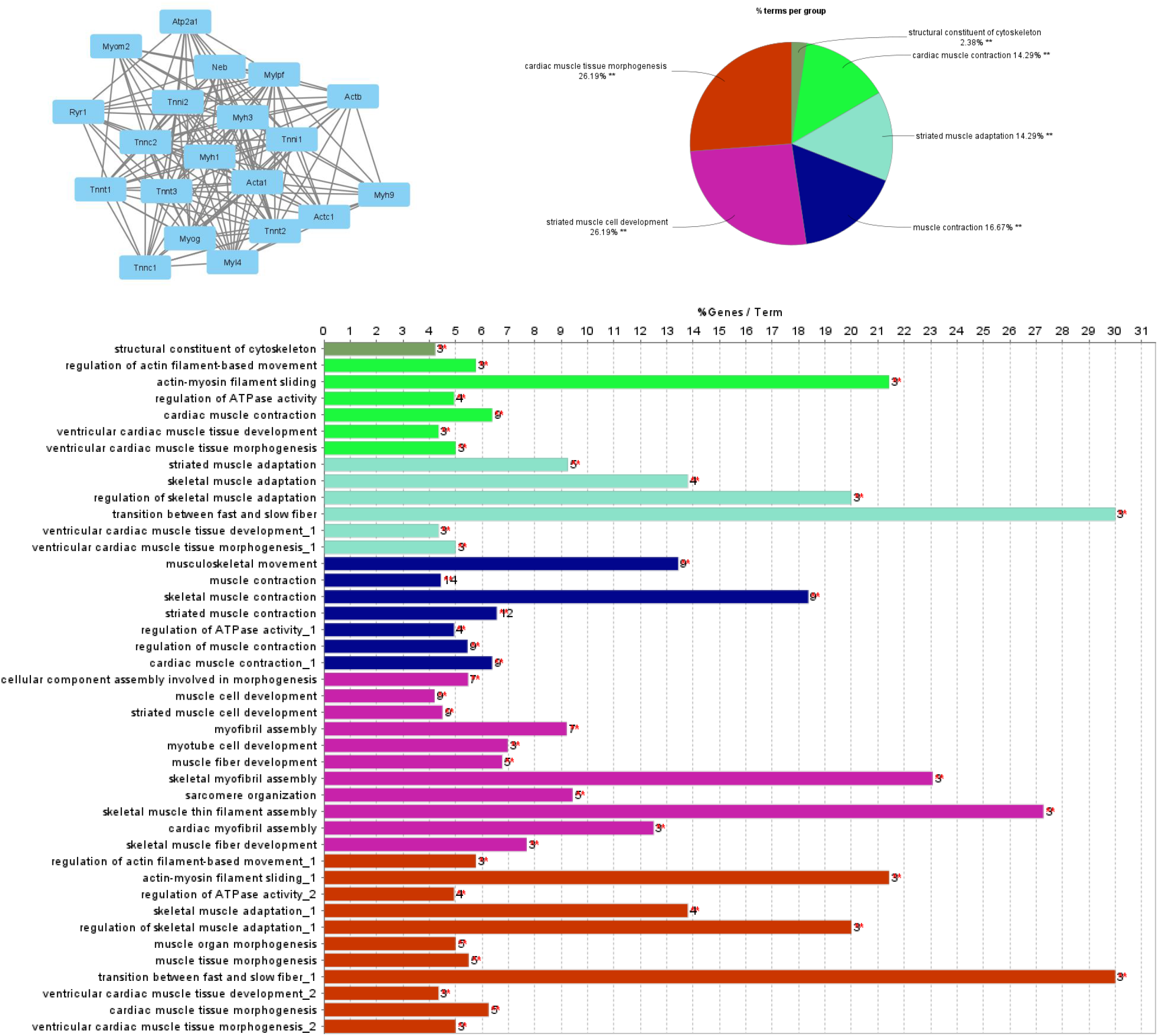
The top module of the protein-protein interaction network and Gene Ontology (GO) terms of DEGs. (A) The top module of the DEGs. (B) Overview chart with GO analysis of biological process in the top module. (C) Functional distribution of GO analysis of biological process in the top.

**Figure 5.**
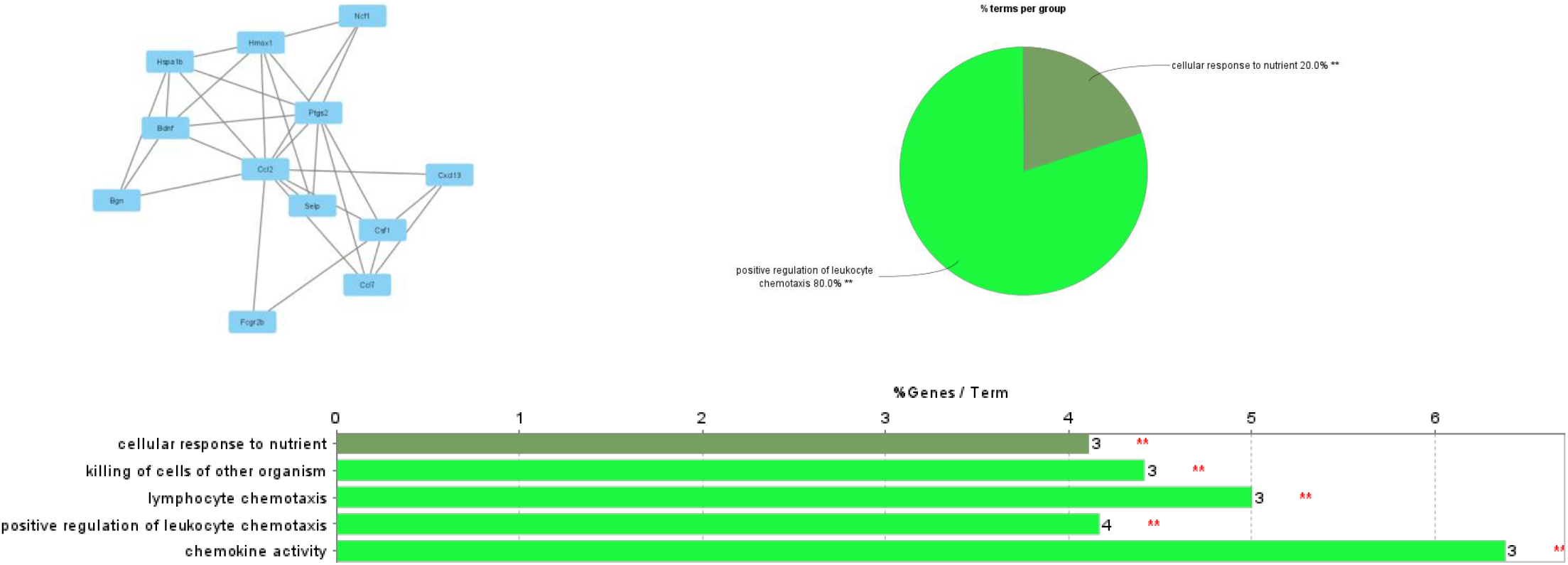
The Second module of the protein-protein interaction network and Gene Ontology (GO) terms of DEGs. (A) The Second module of the DEGs. (B) Overview chart with GO analysis of biological process in the Second module. (C) Functional distribution of GO analysis of biological process in the Second.

## DISCUSSION

In this study, a total of 377 DEG was screened, including 178 upregulated genes and 199 downregulated genes. To gain a deeper understanding of these DEGs, we performed GO and KEGG pathway analysis of these DEGs. GO analysis showed that up-regulated DEG was primarily involved in angiogenesis, cell adhesion, and inflammatory response. We also showed that down-regulated DEG is involved in the regulation of muscle contraction, skeletal muscle fiber development. In addition, the upregulated KEGG pathway of DEG included Rheumatoid arthritis, Chemokine signaling pathway, and Cytokine-cytokine receptor interaction. Downregulated DEG included Calcium signaling pathway, hypertrophic cardiomyopathy (HCM), Dilated cardiomyopathy, Neuroactive ligand-receptor interaction, and Cardiac muscle contraction.

Analysis of two modules selected from the PPI network showed that high-dose pyruvate exposure to C2C12 was primarily associated with muscle contraction, muscle organ morphogenesis, leukocyte chemotaxis, and chemokine activity. Pyruvic acid treatment generally suppresses inflammation (3-6). So far, pyruvate treatment has been reported to inhibit the activation of p38 mitogen-activated protein kinase and NF-κB, two signaling pathways important for cytokine release (5). The results of this study show that high-dose pyruvate treatment reacts differently than low-dose treatment. Increased production of reactive oxygen species after treatment with pyruvate has been observed in many cell lines (12, 13). In previous studies, high concentrations of pyruvate upregulated reactive oxygen species (ROS) through activation of c-jun N-terminal kinase (JNK) in mitochondria (12). In addition, ROS is involved in the inflammatory response (14, 15). Therefore, the chemokines and inflammatory response-related genes found in this study may have been triggered by ROS. Interestingly, muscle contraction and muscle organ morphogenesis were down-regulated. Previous studies have reported that low-dose pyruvate treatment enhances protein synthesis (16). High-dose treatment of pyruvate was found to alter the response of protein metabolism differently than low-dose treatment. All 10 genes identified as hub genes of the PPI network were involved in muscle differentiation. Nine of them were actin, myosin and troponin-related genes. Myog functions as a transcriptional activator that plays a role in muscle differentiation (17, 18). Previous studies have reported that down-regulation of Myog suppresses myocyte differentiation (19). Therefore, the down-regulation of skeletal muscle-related genes in this study may have been caused by a decrease in Myog.

Overall, it is clear that high-doses of pyruvate treatment contribute significantly to genetic alterations. Nevertheless, our research is not without the following limitations: In this study, much more pyruvic acid was added in vitro than the biological standard. Whether the same phenomenon as this research occurs in vivo is the next research topic.

## Conclusion

In summary, our study attempted to reveal molecular changes induced by pyruvate addition in skeletal muscle by bioinformatics analysis. High-dose pyruvate treatment of C2C12 was found to be associated with an increased inflammatory response and decreased skeletal muscle formation. However, further studies are still needed to verify the function of these molecules at high doses of pyruvate.

## Acknowledgements

Not applicable.

